# dGAE(297-391) tau fragment promotes formation of CTE-like full-length tau filaments

**DOI:** 10.1101/2023.02.01.526268

**Authors:** Kristine Kitoka, Alons Lends, Gytis Kucinskas, Anna Lina Bula, Lukas Krasauskas, Vytautas Smirnovas, Monika Zilkova, Branislav Kovacech, Rostislav Skrabana, Jozef Hritz, Kristaps Jaudzems

## Abstract

The microtubule-associated protein tau forms disease-specific filamentous aggregates in several different neurodegenerative diseases. In order to understand how tau undergoes misfolding into a specific filament type and to control this process for drug development purposes, it is crucial to study *in vitro* tau aggregation methods and investigate the structures of the obtained filaments at the atomic level. Here, we used the tau fragment dGAE, which aggregates spontaneously, to seed the formation of full-length tau filaments. The structures of dGAE and full-length tau filaments were investigated by solid-state MAS NMR, showing that dGAE allows propagation of a chronic traumatic encephalopathy (CTE)-like fold to the full-length tau. The obtained filaments efficiently seeded tau aggregation in HEK293T cells. This work demonstrates that *in vitro* preparation of disease-specific types of full-length tau filaments is feasible.

## Introduction

Protein misfolding into insoluble amyloid deposits is a hallmark of many neurodegenerative diseases [1]. The microtubule-associated protein tau is intrinsically disordered and highly soluble [2,3]. Despite this, insoluble tau filaments are formed in several neurodegenerative diseases such as Alzheimer’s disease (AD) [4], corticobasal degeneration (CBD) [5], chronic traumatic encephalopathy (CTE) [6], and Pick’s disease (PiD) [7]. Remarkably, the adopted folds of tau filaments are disease specific [4-8].

In the brain, tau is present as six different isoforms comprising 352-441 residues produced through alternative splicing. The isoforms differ by the number of N-terminal inserts (0N, 1N or 2N) and C-terminal repeats (3R or 4R isoforms) [9]. The filamentous tau structures have been investigated to identify the parts of the protein incorporated into the rigid cores of the filaments [10,11]. Typically, the disease-associated tau filament cores do not exceed one-quarter of the protein sequence. In electron micrographs, the filaments appear to be surrounded by a fuzzy outer coat [12-14]. A recent breakthrough discovery using cryo-EM on patient-derived material revealed that the rigid core of AD paired helical and straight filaments is formed by the V306-F378 fragment [4]. This region largely overlaps with the I297-E391 (dGAE) fragment (Figure 1a), previously found to be the main component of AD patient-derived material [12]. dGAE covers the end of R2, whole R3 and R4 repeats, and the major part of R’ repeat of the full-length (2N4R) tau protein. Although the dGAE fragment is slightly longer than the AD tau filament core identified by cryo-EM [4], Lovestam et al. showed that the residues at its N- and C-termini are essential for filament formation *in vitro* because several shorter tau constructs were ineffective in forming filaments [15]. Additionally, they proved by high-throughput cryo-EM structure determination that dGAE can form filaments *in vitro*, resembling those found in the CTE and AD patient brains [15,16].

**Figure 1.**
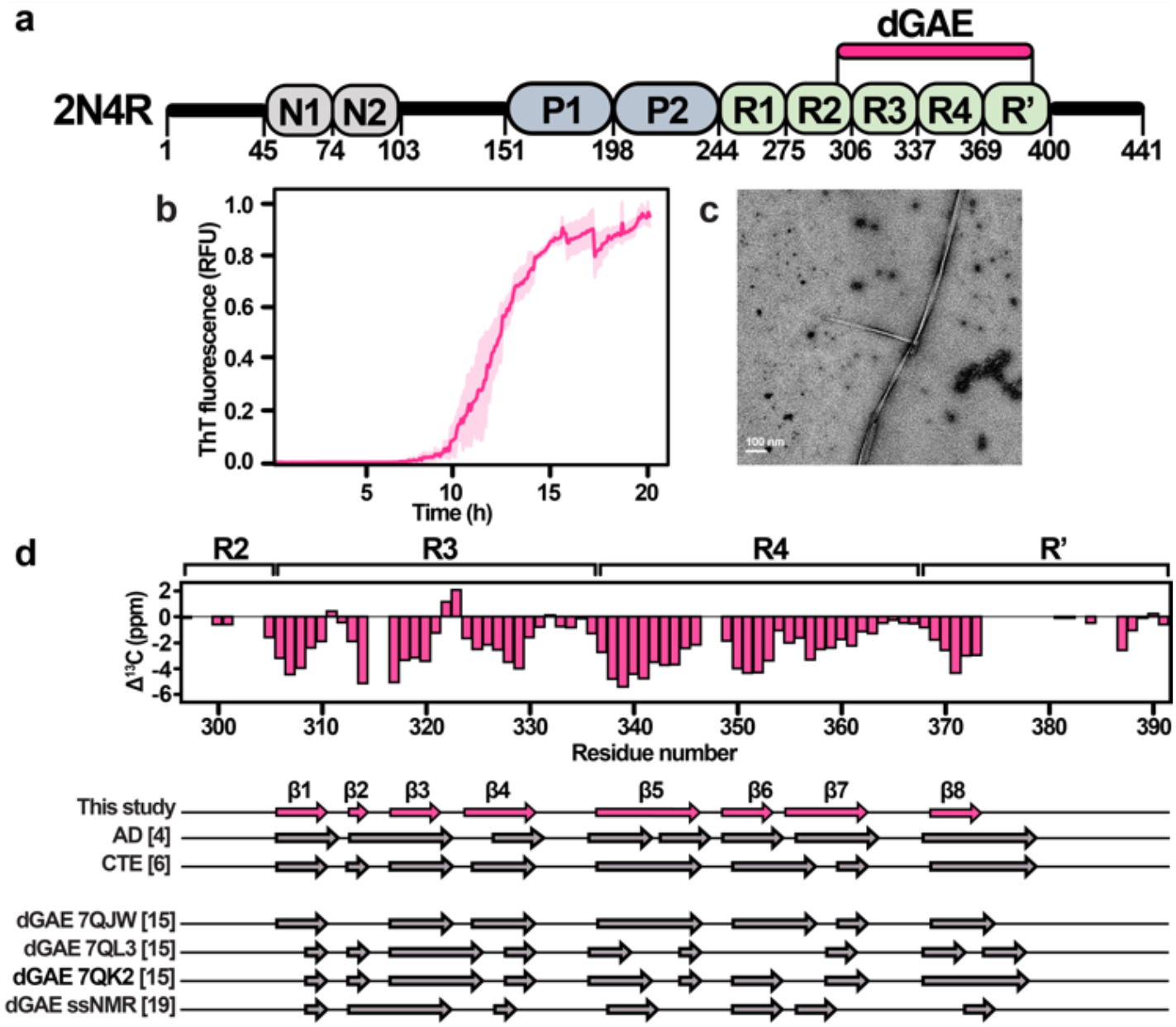
(a) Full-length tau 2N4R domain architecture. N1 and N2 are exon-coded inserts, P1 and P2 indicate the proline-rich regions, and R1 to R’ indicate repeat domains. dGAE includes residues I297-E391 from the repeat domains of the 2N4R full-length tau. (b) ThT fluorescence curve of dGAE aggregation kinetics (average of three replicates with standard deviation values). (c) Negative-stain EM micrograph of dGAE(297-391) tau filaments. (d) Comparison of secondary structures between dGAE filaments presented here, patient-derived tau filaments [4,6], and other *in vitro* dGAE studies [15,19].

In contrast to full-length tau, dGAE aggregates in the absence of any aggregation inducers, allowing a preparation of filaments *in vitro* without heparin [17-19]. Heparin screens electrostatic interactions, thereby inducing conformational rearrangement of the tau protein that leads to its self-assembly [20,21]. Until recently, heparin was the most used cofactor for preparing tau filaments, as it ensures relatively high aggregation yields [22-27]. However, cryo-EM has shown that heparin-induced tau filaments are structurally heterogeneous and distinct from the patient-derived filaments, raising questions about the relevance of such aggregation protocols [28]. Thus, dGAE provides a more biologically relevant route to generate tau filaments for *in vitro* studies, necessary for the development of new disease-modifying therapies for tauopathies. However, it remains elusive whether the findings from dGAE studies can be directly translated into the context of the full-length tau.

Here, we use the solid-state NMR (ssNMR) and other complementary biophysical methods to characterize the structures of both the truncated tau fragment dGAE and the full-length tau filaments obtained by seeding with dGAE. By comparing the spectra of dGAE and seeded full-length 2N4R filaments, we demonstrate that this approach allows the preparation of CTE-like full-length tau filaments *in vitro*. Additionally, the filaments efficiently induced tau aggregation in the FRET biosensor cell line.

## Results

dGAE filaments were formed by incubating 400 μM protein solution with agitation (800 rpm) without cofactors in PBS pH 7.4 and in the presence of 5 mM DTT. Thioflavin T (ThT) fluorescence showed a typical sigmoidal curve indicative of amyloid formation via primary nucleation, elongation, and secondary nucleation (Figure 1b) [29]. The average lag time for filament formation was approximately 8 h, after which an exponential increase in ThT fluorescence was observed with a half-time of 12 h, reaching a plateau at 20 h. The small standard deviation values indicate repeatable aggregation kinetics of dGAE under these conditions. SDS-PAGE analysis (Figure S1) confirms that most of dGAE was aggregated at the end of the incubation period. The electron micrographs show the formation of slightly twisted and unbranched filaments (Figure 1c).

The ssNMR was performed on uniformly ^13^C,^15^N-labeled dGAE filaments using ^13^C-detected experiments at 12 kHz MAS. The 2D DARR ^13^C-^13^C correlation spectrum with 20 ms mixing time displayed line widths of 0.6 ppm for well-isolated peaks, suggesting that the filament preparation yielded a single conformation. Examination of several residue-type-specific regions indicated the expected number of correlations (Figure S2), corroborating high homogeneity of the sample.

Sequence-specific assignments of the backbone ^15^N and/or ^13^C chemical shifts were obtained for residues V306-D314, V318-F346, R349-K369, I371-T373, and D387-E391 from the 3D NCACX, NCOCX and CANCO experiments (Figure S3). Side chain assignments were completed using the 2D DARR spectra with longer mixing times. To simplify the spectra and confirm the assignments, selectively unlabeled samples were prepared by suppressing either lysine or leucine, isoleucine, valine, and lysine labeling (Figure S4). The obtained assignment suggests that at least 60 residues spanning the region V306-T373 are incorporated into the rigid core of our dGAE filaments (Figure S5A). The chemical shifts of C322 correspond to a reduced cysteine, in agreement with the reducing conditions used for the aggregation (Figure S2A). Residues D387-E391 were assigned, implying that the C-terminal part folds back and attaches to the filament core.

To probe the mobile parts outside the rigid filament core, we recorded INEPT-based experiments that allow the detection of highly dynamic residues [30]. In the INEPT 2D 1H-13C correlation spectrum (Figure S5B), the cross peaks exhibiting random coil chemical shifts were assigned to methionine, isoleucine, lysine, histidine, valine, glycine, alanine, phenylalanine, proline, serine, leucine, arginine, and threonine residues. Based on the uniqueness of some amino acid types outside the assigned core, these mobile residues most likely belong to fragments I297-S305 and H374-T386. Altogether the MAS ssNMR data indicate that the dGAE rigid core is composed of residues V306-T373, which interacts with the very C-terminus (residues D387-E391). The N-terminus (residues I297-S305) and residues H374-T386 near the C-terminus are mobile, whereas the rest are semi-rigid or conformationally heterogeneous as they are not observed in either cross-polarization (CP) or INEPT-based experiments.

To identify the locations of secondary structures and make a comparison with other tau structures, deviations of backbone Cα and Cβ chemical shifts from random-coil values, known as the secondary chemical shifts (SCS), were analyzed. Most of the assigned residues were found to adopt a β-sheet conformation, whereas unassigned fragments coincide with loops in the cryo-EM structures (Figure 1d). Strand β1 covers the R3 hexapeptide motif V306–Y310, β2 is formed by V313-D314, β3 by K317-K321, and β4 by S324-H330. V337-F346 strand β5 and covers the R4 hexapeptide motif V337-E342, R349-K353 forms β6, and L355-V363 forms β7. K369-T373 forms β8 at the beginning of R’. As expected, both hexapeptides are involved in dGAE filament formation. Compared to the cryo-EM structures of patient-derived tau, our assignment covers residues that belong to β1-β8 strands of the cryo-EM structures. The locations of the identified β strands fit best to the CTE protofilament cryo-EM data (Figure 1d). We also compared the assigned β-strand locations with other dGAE *in vitro* studies, performed by cryo-EM (PDB: 7QJW, 7QL3, 7QK2) [15] and by ssNMR [19]. The comparison shows that the β-strand locations differ notably, especially in the region of R4 hexapeptide. The 7QJW structure, which is very close to the CTE type II filaments [15], fits best with our data.

The AD and CTE protofilament structures are both C-shaped, with the CTE protofilament fold adopting a more extended conformation. The subtle conformational differences yield alternative side chain packing arrangements resulting in a distinct cross-peak pattern between V337, V339, and I354 and L325, I328, and V363 (Figure 2a,c). To confirm the structural similarity of the analyzed dGAE filaments to a disease-specific filament type, we examined the dipolar recoupling DARR spectra with long mixing times. Two cross-peaks belonging to V339Cβ-I354Cδ1 and V337Cβ-I354Cγ2 resonances were identified, reporting on the packing of the I354, V337, and V339 residues (Figure 2a). This indicates that the side chain of I354 is packed between V337 and V339. Based on the available cryo-EM structures, such a cross-peak pattern agrees well with the CTE β-helix arrangement (Figure 2b right panel) and not the AD PHF-like β-helix arrangement (Figure 2b left panel). Furthermore, several cross-peaks between L325, I328, and V363 were identified (Figure 2c), suggesting that V363 is packed between L325 and I328. Such an arrangement is consistent with the CTE protofilament packing (Figure 2d left panel) but not with the packing in the 7QL3 and 7QK2 dGAE structures (Figure 2d middle and right panel), where L325 and I328 are solvent-exposed. By examining DARR spectrum with 500 ms mixing time, we found cross-peak patterns corresponding to inter-protofilament side chain contacts of H329 and E338 (Figure 2e). This unambiguous long-range contact indicates that intermolecular contacts involve ^332^PGGG^335^ motifs facing each other in an anti-parallel manner. This arrangement places H329 and E338 near one another, which is consistent with the CTE type II fold (Figure 2f).

**Figure 1.**
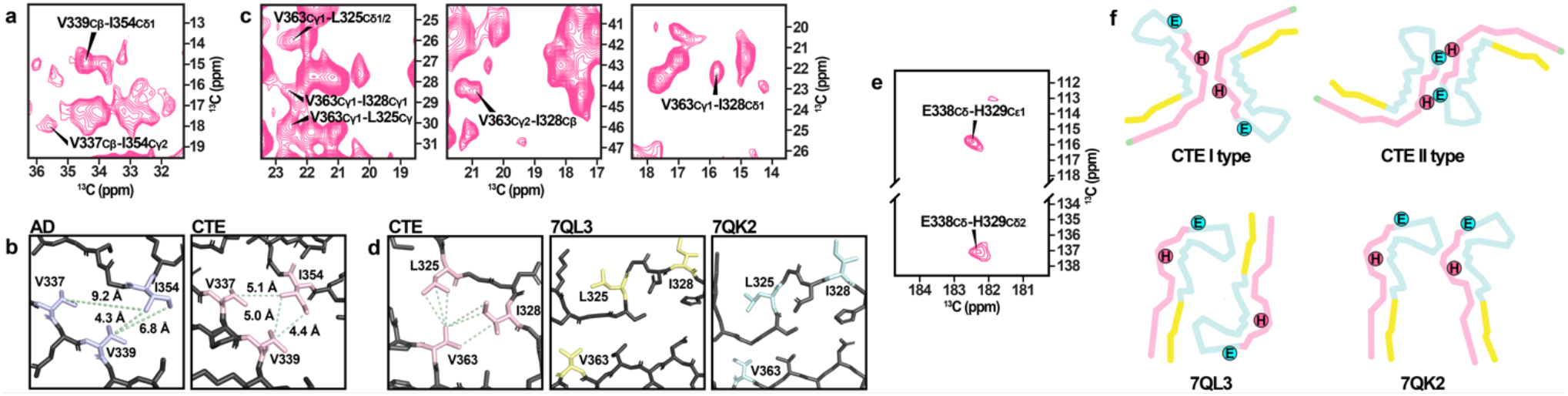
(a) A selected region of a 2D ^13^C-^13^C DARR spectrum (150 ms mixing time) showing V339Cβ-I354Cδ1 and V337Cβ-I354Cγ2 correlations. (b) V337, V339, and I354 side-chain packing in AD and CTE filaments. Selected regions of a ^13^C-^13^C DARR spectrum (150 ms mixing time) showing V363Cγ1-L325Cδ1/2, V363Cγ1-I328Cγ1, V363Cγ1-L325Cγ, V363Cγ2-I328Cβ, and V363Cγ1-I328Cδ1 correlations. (d) L325, I328, and V363 side-chain packing in CTE, 7QL3, and 7QK2 filaments. (e) Selected regions of a ^13^C-^13^C DARR spectrum (500 ms mixing time) showing E338Cδ-H329Cε1 and E338Cδ-H329Cδ2 correlations. (f) H329 and E338 locations in the CTE, 7QL3, and 7QK2 filament structure.

The obtained dGAE filaments were used as a template for seeded aggregation of full-length tau, using identical conditions as for dGAE alone. The efficiency of aggregation was assessed by SDS-PAGE (Figure 3a), which showed that a major part of the full-length tau protein remained in the solution (Figure 3a, lane S), and only a minor part was in the pellet (Figure 3a, lane P). This suggests that full-length tau reaches an equilibrium of aggregation/dissolution processes even in the presence of dGAE seeds. Template-driven aggregation was also monitored using ThT fluorescence (Figure 3b). In contrast to the dGAE fluorescence curve, the dGAE seeded 2N4R aggregation lacked the lag phase and was dominated by elongation (compare with Figure 1b). This shows that dGAE filaments can template full-length tau even when the nucleation of 2N4R alone is unfavorable. The obtained full-length tau filaments were long and formed a net-like arrangement on EM grids (Figure 3c). Compared with dGAE filaments, the body of the 2N4R filaments seems to be covered by additional densities, most likely the fuzzy coat. In the AFM micrographs, the 2N4R filaments appear to be twisted (Figure S6).

**Figure 3.**
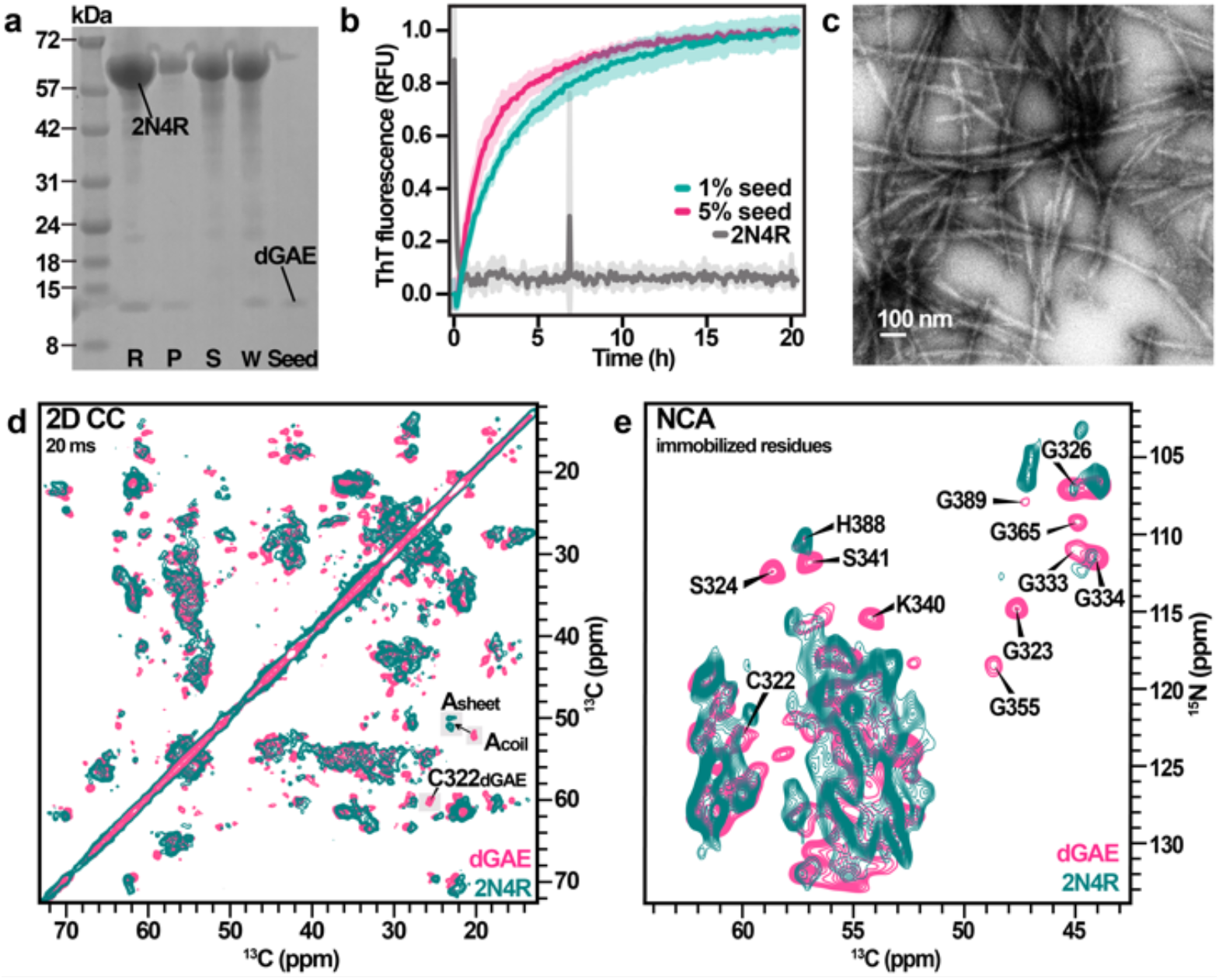
(a) SDS-PAGE of the full-length tau sample with dGAE seeds before and after aggregation in PBS pH 7.4, 5 mM DTT. Lane R is a reference sample at the beginning of aggregation, lane P corresponds to the pellet fraction after aggregation, lane S – the supernatant fraction after aggregation, lane W -whole reaction mixture after aggregation, and lane Seed is the dGAE seed. (b) ThT fluorescence curve of dGAE-seeded 2N4R aggregation kinetics (average of four replicates with standard deviation values). (c) Negative-stain EM micrograph of the seeded full-length tau filaments. Overlay of 2D ^13^C-^13^C DARR spectra of dGAE and seeded full-length tau filaments. (e) Overlay of 2D NCA spectra of dGAE and seeded full-length tau filaments. The missing residues are identified

Structural analysis of dGAE-seeded full-length tau filaments was performed by comparison of 2D ssNMR spectral fingerprints with dGAE filaments. Full-length tau filaments exhibit more signals in the ^1^H-^13^C INEPT spectrum compared to dGAE (Figure S7), confirming the formation of full-length tau filaments with mobile flanking regions. Several resonances, such as isoleucine and alanine, overlap in these spectra, and some become more intense in the spectrum of full-length tau filaments.

The DARR spectra of dGAE and full-length tau filaments look quite similar. However, notable differences are observed in a few spectral regions (Figure 3d). Alanine Cα, Cβ resonances, which in the DARR spectrum of dGAE exhibit random coil chemical shifts, are shifted toward beta-sheet chemical shifts. The increased number of alanine resonances and the fact that one of these resonances most likely belongs to A390 suggests that additional β-strands outside V306-F378 are present the full-length tau filaments. This leads us to think that the C-terminal part also contributes to the 2N4R filament rigid core. In particular, this is an interesting observation because these parts were not detected in cryo-EM studies of CTE tau filaments. Furthermore, the C322 peak is significantly shifted/missing in the DARR spectrum of full-length tau filaments, indicating an altered C322 conformation and/or dynamics.

The overlay of 2D NCA spectra (Figure 3e) shows the disappearance or shifting of several dGAE resonances in the full-length tau spectrum. These include the C322-S324 fragment and several glycine residues. Surprisingly, both K340 and S341 also disappear, although they are part of the R4 hexapeptide V337-E342. At the same time, the INEPT spectrum of 2N4R (Figure S7) does not exhibit any new resonances that could belong to serine. These changes suggest that the R4 hexapeptide is influenced by the mobile regions of full-length tau filaments when incorporated into full-length filaments.

In order to verify the dGAE and dGAE-seeded full-length tau filament seeding potential in cells, we tested their proteopathic seeding activity on Tau RD P301S FRET biosensor epithelial cell line [31] (Figure 4a). Both filament types consistently exhibited potent seeding activity in the cells. The averaged FRET fluorescence signal was higher for dGAE-seeded 2N4R filaments than for dGAE filaments (Figure 4b), reflecting a greater extent of per-cell aggregation of the intracellular reporter tau after transfection with the 2N4R filaments compared to transfection with the dGAE filaments. This may be due to the presence of additional β-strands in the 2N4R filaments, as suggested by the DARR spectra (Figure 3d), which confer a larger surface for seeded aggregation of the biosensor reporter tau.

**Figure 4.**
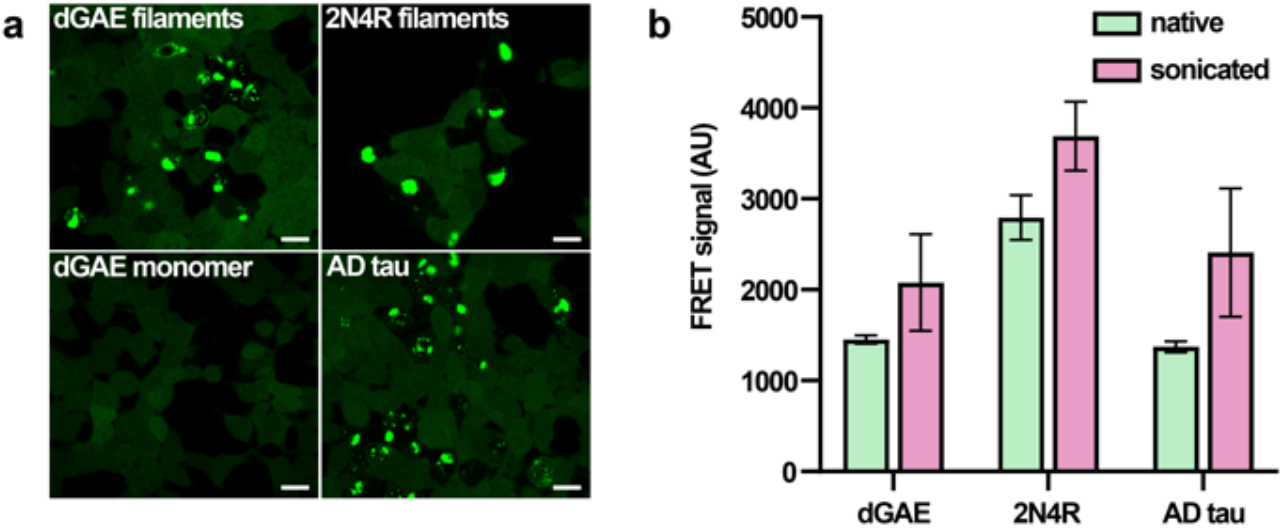
Tau aggregation in biosensor cell line induced by dGAE and 2N4R filaments. (a) Fluorescence microscopy of tau inclusions after transfection with filaments. dGAE monomers and sarkosyl insoluble AD tau filaments were used as negative and positive control, respectively. White bars represent 20 μm. (b) Quantification of intracellular tau inclusions in FRET-positive cells based on averaged median fluorescence intensity in the FRET channel. Error bars represent standard deviation.

## Discussion

Recently, several groups have studied filaments formed by the dGAE fragment [15,16,19,32-34]. Lovestam et al. showed that the use of NaCl or PBS gives rise to CTE type II filaments or new morphologies [15,16]. Based on the secondary structure data (Figure 1c), β-strand locations in our obtained dGAE filaments fit the best with CTE protofilament β-strand locations, except that the β8 strand was not observed either in the CP or INEPT type spectra. Although the solution conditions were quite similar, we did not observe any resonances that correspond to minor conformations and different β-strand location patterns. These differences could be attributed to different shaking speeds and aggregation volumes. Our determined β-strand locations also differ from another ssNMR study of dGAE filaments formed under different aggregation conditions (10 mM NaPi pH 7.4, 10 mM DTT). Interestingly, the rigid core identified in this study also lacked most of the C-terminal beta-strand from the cryo-EM structures [19].

Our obtained dGAE filaments adopt the beta-helix conformation seen in CTE-type protofilaments, suggesting that disease-specific CTE type II filaments can be generated *in vitro*. Usually, type II is a minor fold found in the CTE brain. However, increased type II levels have been found in amyotrophic lateral sclerosis/parkinsonism dementia (ALS/PDC) complex [35] and subacute sclerosing panencephalitis (SSPE) [36], suggesting that similar molecular mechanisms underlie these diseases. Inflammation processes may be a shared feature among CTE, SSPE, and ALS/PDC. To date, the major bottleneck in the structural studies of full-length tau filaments is the high solubility of tau. Spontaneous tau aggregation without cofactors is usually less efficient than cofactor-induced aggregation. In addition, aggregation becomes even less efficient by increasing the length of tau from truncated constructs to full-length isoforms [37,38]. This study shows that the full-length tau protein can be efficiently aggregated by template-driven aggregation using the truncated tau fragment dGAE. The observed elongation profiles suggest that the dGAE filaments propagate their fold onto the full-length 2N4R tau, which may pave the way for disease-specific full-length tau filament studies.

Both dGAE and seeded 2N4R filaments efficiently induced tau aggregation in the Tau RD P301S FRET biosensor cell line. However, it remains to be clarified if a similar mechanism exists for the tau aggregation in the brain, where “tauons” generated by tau truncation have been proposed to induce a template-assisted pathology propagation [39].

Tau neurofibrillary tangles are known to be hyperphosphorylated. However, the established phosphorylation sites in CTE are located outside the dGAE sequence [40]. Thus, dGAE aggregation should not be affected by phosphorylation. On the other hand, seeded aggregation of full-length tau could be affected by hyperphosphorylation. Further studies to compare phosphorylated dGAE-seeded full-length tau with wild-type tau are warranted.

In summary, we used ssNMR spectroscopy to investigate the features of full-length and truncated tau filaments generated from the dGAE fragment. The rigid core of the obtained single-conformation dGAE filaments spans residues V306-T373 covering residues up to the β8 strand seen in cryo-EM structures. Inter-nuclear contacts indicate that the filament tertiary and quaternary structure corresponds to that found in CTE patient-derived type II filaments. The dGAE CTE type II fold can be propagated to full-length tau, and both types of filaments efficiently seed tau aggregation in cells. This work sets the basis for future ssNMR spectroscopy studies of disease-specific tau filaments. In particular, a deeper understanding of the interactions between the rigid core and the mobile flanking regions could help to identify important drug-binding sites, facilitating the design of more effective therapeutic candidates against the progression of AD and other tauopathies.

## Acknowledgements

We acknowledge funding from the Latvian Council of Science grant no. lzp-2019/1-0244 and European Union’s Horizon 2020 research and innovation programme under the Marie Skłodowska-Curie Rise InterTau grant agreement No 873127. K.K. is supported by “Mikrotīkls doctoral scholarship in the field of exact and medical sciences”, administered by the University of Latvia Foundation. A.L. is supported by Marie Curie widening fellowship nr. 101038074—”Oligomers-MAS-NMR”. G.K. and J.H. acknowledge the Czech Science Foundation (no. GF20-05789L) and Nanobiotechnology and Cryo-electron microscopy and tomography core facilities CEITEC MU of CIISB, Instruct-CZ Centre supported by MEYS CR (LM2018127). R.S. is supported by the Slovak Research and Development Agency grant APVV-21-0479 and VEGA grant 2/0141/23, M.Z. is supported by VEGA grant 2/0134/22 and B.K. is supported by grant APVV-20-0585.

