## Supporting information for "dGAE(297-391) tau fragment promotes formation of CTE-like full-length tau filaments"

### Methods

**Protein expression.** The gene encoding dGAE(297-391) cloned into a pET-17b vector was transfected into *E. coli.* BL21(DE3) competent cells. A starter culture was grown in 50 mL LB medium containing 100 µg/mL ampicillin. After overnight growth at 30 °C with 200 rpm shaking, 20 mL of the overnight culture were harvested and resuspended into 1 L LB medium containing 100 µg/mL ampicillin. Cells were grown at 37 °C and 200 rpm shaking until $OD_{600}$ reached ~0.8. At this point, cells were spun down at 7,000 g and 4 °C for 15 min. The cell pellet was resuspended in 500 mL M9 minimal media (1 g/L $^{15}NH_4Cl$, 2 g/L D-Glucose-$^{13}$C6, Cambridge Isotope Laboratories, Inc.), 2 mM $MgSO_4$, 0.1 mM $CaCl_2$, vitamin/mineral supplements, and 100 µg/mL ampicillin). For selectively unlabeled samples, either 400 mg/L unlabeled lysine or 230 mg/L unlabeled valine, 230 mg/L unlabeled isoleucine, 230 mg/L unlabeled leucine, and 400 mg/L unlabeled lysine[1,2] were added to M9 minimal media. Cells were then grown at 37 °C and 200 rpm shaking for 60 min before induction with IPTG (0.1 mM). Cells were harvested 3 h after induction by centrifugation at 7,000 g and 4 °C for 15 min. The cell pellets were resuspended in 10x 50 mM NaPi pH 6.9 buffer and stored at -80 °C. The gene encoding 2N4R full-length tau was cloned into a pET-28a vector. Uniformly $^{13}$C-$^{15}$N labeled full-length 2N4R tau was expressed as described above.

**dGAE purification.** The cell pellet in lysis buffer (50 mM NaPi pH 6.9, 8 M urea, 1 mM EDTA, 2 mM DTT, 0.1 mM PMSF) was sonicated on ice (1 s on, 4 s off, 15 min) using a probe sonicator. Sedimentation of insoluble particles was done by centrifugation at 40,000 g for 45 min. The supernatant was dialyzed overnight at 4 °C against buffer A (50 mM NaPi pH 6.9, 1 mM EDTA, 2 mM DTT, 0.1 mM PMSF) to remove urea. The following day, the sample was filtered and applied onto an equilibrated cation exchange chromatography column (HiTrap SPHP, GE Healthcare). Elution was done using a linear gradient of 0-40 % of buffer B (50 mM NaPi pH 6.9, 1 M NaCl, 1 mM EDTA, 2 mM DTT, 0.1 mM PMSF) in 3 column volumes (CV) and a linear gradient of 40-100 % of buffer B in 1 CV. Fractions in the range of eluate conductivity 7–40 mS/cm were collected. The presence of protein was confirmed using SDS-PAGE. Fractions containing dGAE were pooled, and excess Milli-Q water was added to reach the conductivity of 5-6 mS/cm of the pooled sample. The solution was centrifuged at 20,000 g for 10 min at 4 °C. The supernatant was filtered and applied onto an equilibrated cation exchange chromatography column (HiTrap SPHP, GE Healthcare) for the second cation exchange. Elution was done using a linear gradient of 0-30 % buffer B in 10 CV and a linear gradient of 30-100 % buffer B in 1 CV. Fractions containing dGAE were purified by size-exclusion chromatography (HiLoad Superdex 75 26/60 column, GE Healthcare) in PBS pH 7.4, 5 mM DTT. Purified tau protein was stored in PBS pH 7.4, 5 mM DTT at −80 °C for further experiments. The concentration of dGAE was estimated using a sequence-specific extinction coefficient for absorbance at 280 nm. The expression and purification protocol yields ~30 mg uniformly $^{13}$C-$^{15}$N labeled dGAE per 1 L of M9 minimal media.

**2N4R purification.** The cell pellet in lysis buffer (50 mM NaPi pH 7.4, 8 M urea, 20 mM imidazole, 500 mM NaCl, 1 mM EDTA, 0.1 mM PMSF) was sonicated on ice (1 s on, 4 s off, 15 min) using a probe sonicator. Sedimentation of insoluble particles was done by centrifugation at 40,000 g for 45 min. The supernatant was dialyzed overnight at 4 °C against buffer A (50 mM NaPi pH 7.4, 20 mM imidazole, 500 mM NaCl, 1 mM EDTA, 0.1 mM PMSF) to remove urea. The following day, the sample was filtered and applied onto an equilibrated immobilized metal affinity chromatography column (HisTrap HP, GE Healthcare). Elution was done by applying a linear gradient of 0-100 % of buffer B (50 mM NaPi pH 7.4, 500 mM imidazole, 500 mM NaCl, 1 mM EDTA, 0.1 mM PMSF) in 10 CV. Fractions containing 2N4R were pooled and dialyzed against TEV protease cleavage buffer (50 mM Tris pH 8, 150 mM NaCl, 1 mM EDTA, 2 mM DTT). The removal of the His-tag was done by overnight TEV protease cleavage (1:10 molar ratio TEV:2N4R) in dialysis at 4 °C. The next day, the cleavage mixture was heated at 75 °C for 15 min before being centrifuged at 20,000 g and 4 °C for 20 min. The supernatant was purified by size-exclusion chromatography (HiLoad Superdex 200 26/60 column, GE Healthcare) in PBS pH 7.4, 10 mM DTT to achieve the maximal purity of 2N4R. Purified tau protein was stored in PBS pH 7.4, 10 mM DTT at −80 °C for further experiments. The concentration of 2N4R was estimated using a sequence-specific extinction coefficient for absorbance at 280 nm. The expression and purification protocol yields ~20 mg uniformly $^{13}$C-$^{15}$N labeled 2N4R per 1 L of M9 minimal media.

**dGAE filament assembly.** Before aggregation, the dGAE solution was buffer-exchanged to fresh PBS pH 7.4, 5 mM DTT. 400 µM $^{13}$C-$^{15}$N labeled dGAE was distributed into 1.5 mL microcentrifuge tubes with a final volume of 1 mL per tube and agitated at 37 °C and 800 rpm orbital shaking with 2 mm orbit (Thermo–Shaker TS-100C, Biosan) for 48 h. The aggregation of the tau proteins was verified by SDS-PAGE.

**2N4R seeding.** To seed full-length 2N4R tau, dGAE filaments were sonicated for 2 min in a water bath. 1 % (w/w) of seeds were added to 100 µM monomeric 2N4R tau in the PBS pH 7.4, 5 mM DTT. Seeded 2N4R aggregation was done under the conditions used for dGAE aggregation. The samples were shaken for ~96 h.

**ThT fluorescence.** To monitor aggregation kinetics, Thioflavin T (ThT) was added to the protein at a final concentration of 100 µM. 100 µL of sample per well were pipetted in a 96 well plate (Half Area Plate, Corning). ThT fluorescence was measured with an excitation at 440 nm with an excitation bandwidth of 10 nm and emission at 480 nm with a bandwidth of 10 nm (Clariostar Plus plate reader, BMG Labtech). The aggregation assay was performed at 37 °C with 700 rpm orbital shaking. For seeded aggregation experiments, dGAE seeds formed using Thermo–Shaker TS-100C shaker were used (described in *dGAE filament assembly*). Measurements were read at an interval of 5 min. Each sample type was measured triplicates (dGAE) or quadruplicates (dGAE seeded 2N4R aggregation). The analysis of the aggregation data was performed using Amylofit software (http://www.amylofit.ch.cam.ac.uk)[3].

**Negative-stain electron microscopy.** The filaments were centrifuged at 16,000 g for 20 min. The pellets were separated and resuspended in 50 µL of PBS pH 7.4, 5 mM DTT. Due to the high concentration of filaments, a sample was diluted 100 times with an aggregation buffer (PBS pH 7.4, 5 mM DTT). Copper grids with a layer of 12 nm carbon foil were cleaned for 15 s using hydrogen plus oxygen plasma glow discharger Gatan Solarus II (Gatan–Ametek). 4 µL of the sample was applied on the grid for 1 min and stained by 16 µL of 2 % uranyl acetate. Visualizations were done by Talos F200C transmission emission microscope equipped with Ceta-D camera (ThermoFisher). Micrographs were collected at 45k magnification with a pixel size of 3.23 Å and a dose rate of 35 electrons per $Å^2$.

**Atomic force microscopy.** Filaments were immobilized on clean home-made mica plates and incubated for 10 min. After the incubation period mica was cleaned for 4-5 times with a rough flow from pipette with 200 µL of Milli-Q water. Washed mica was placed into a vacuum chamber for 30-35 min to dry. Dried mica was placed into a vacuum fixed holder of Bruker Dimension FastScan atomic force microscope (Bruker). Microscope was equipped with the Bruker SAA-HPI-SS probe (Bruker). Peak Force set point was 500 pN, scanning speed 0.8 Hz, rest of parameters were automatically set up by ScanAsyst module. Data was processed using NanoScope Analysis ver.2.0. Final AFM images were processed using the Gwyddion software package.

**MAS NMR rotor packing.** Aggregated samples were pooled together and spun at 20,000 g and 4 °C for 30 min. The supernatant was then removed. The pellets were packed into 3.2 mm MAS rotors (Bruker) using a rotor packing device (Giotto Biotech) by centrifugation at 30,000 g.

**MAS NMR Experiments.** Spectra for assignment of $^{13}C$ and $^{15}N$ resonances were acquired on an 800 MHz Bruker Avance III HD spectrometer equipped with a 3.2 mm H/C/N E-free MAS probe. The MAS frequency for experiments was 12 kHz. The temperature regulation was set to 273 K using a Bruker BCU II cooling unit. 3D spectra were measured in blocks and co-added. A full list of experimental details of the used acquisition and processing parameters used for the NMR experiments recorded are given in Table S1 for dGAE samples and in Table S2 for dGAE-seeded full-length 2N4R tau sample.

**Analysis of the solid-state NMR data.** All spectra were processed with Bruker Topspin software. Chemical shift assignment was done using in CcpNmr AnalysisAssign[4]. The assignment was done by using resolved and unique resonances such as Thr, Pro, Cys, Ser, Tyr, etc. as starting points for finding connectivities for sequential assignment by means of NCACX, NCOCX, and CANCO experiments. DARR with mixing times of 100, 150, 200, and 500 ms were used to obtain intraresidue, sequential, medium- and long-range correlations. $^{13}C\alpha$ and $^{13}C\beta$ chemical shift deviations from random-coil values[5] were used to determine secondary chemical shifts values. The secondary chemical shifts calculated using the $\Delta\delta C\alpha$-$\Delta\delta C\beta$ equation, the value for each residue was determined as an average over three residues. Assigned chemical shifts have been deposited in the Biological Magnetic Resonance Data Bank under accession code 52070.

**Liposome-mediated transduction of tau seeds.** dGAE filaments and 2N4R filaments seeded by dGAE were identical to filaments used in the ssNMR experiments. Sarkosyl insoluble AD tau filaments for positive control were prepared as previously described[6]. Filament samples were used without further treatment or after sonication in a glass tube (glass insert 250 µL, CHS-INS-63, Chromservis SK) in a water bath for 10 min. dGAE was precleared by centrifugation at 100,000 g for 2 h at room temperature and supernatant containing monomeric tau was used as a negative control. Tau RD P301S FRET Biosensor cells (ATCC® CRL-3275™), prepared by transducing HEK293T cells with 2 separate lentivirus constructs encoding tau RD P301S-CFP and tau RD P301S-YFP[7], were seeded at 60% confluency on 24-well plates a day before the experiment. Cells were transduced with tau samples using Lipofectamine™3000 reagent (Invitrogen).Transduction complexes were prepared by mixing 25 µL Opti-MEM (Gibco) with 1 µL Lipofectamine™3000 and proteopathic tau samples diluted in OptiMEM in a total volume of 52 µL per well. Tau-lipid complexes were incubated at room temperature for 15 min, added to the cells, which were then incubated for further 24 h before flow cytometry analysis.

**Fluorescence Resonance Energy Transfer (FRET) Flow cytometry.** Transduced cells were collected and the BD LSRFortessa™ II cell analyzer was used for FRET flow cytometry. To measure CFP and FRET, cells were excited with the 405 nm laser, and fluorescence was monitored with a 450/40 nm and 525/50 nm filter, respectively. To measure YFP, cells were excited with a 488 laser and fluorescence was monitored with a 525/50 nm filter. To select FRET-positive cells and quantify FRET, we used a gating strategy and evaluation as previously described[6]. Tau seeding activity was presented as an average of median fluorescence intensity (MDI) of FRET-positive cells, using at least two independent cell cultures. MDI was evaluated from 1000 FRET-positive cells.

**Fluorescence microscopy.** Tau biosensor cells were plated on Nunc Lab-Tek II Chamber Slide System (Merck) and transduced with tau samples. Representative live cell images of YFP fluorescence were captured by LSM 710 confocal microscope (Zeiss, Jena, Germany) five days after transduction.

### Results and Discussion

**Figure S1.** SDS-PAGE of dGAE before and after aggregation in PBS pH 7.4, 5 mM DTT. Lane P corresponds to the pellet fraction after aggregation, lane S – the supernatant fraction after aggregation shows two distinct bands corresponding to dGAE and a truncated product left after the purification process, and lane R – a reference sample before dGAE aggregation. dGAE assembled into filaments almost quantitatively.

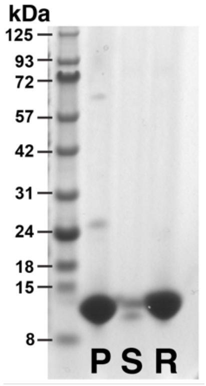

**Figure S2.** (a) DARR spectrum of U-$^{13}$C,$^{15}$N-labeled dGAE filaments at 12 kHz MAS. A single C322 cross-peak suggests that cysteine in dGAE filaments adopts single conformation as well as the residues close to C322. At least five correlation peaks are observed in the serine Cα-Cβ range, which is in good agreement with the presence of six serine residues within the dGAE rigid core. In the case of threonine residues, several Cβ-Cγ cross-peaks are observed, while the Cα-Cβ region exhibits one very intensive cross-peak. In the proline Cβ-Cδ and Cγ-Cδ region, at least three pairs of resonances are seen, which is in line with three proline residues within dGAE rigid core. The CC spectrum displays five well-separated isoleucine resonances in the Cβ-Cγ2 region. Moreover, three display strong Cγ1-Cγ2 signals. Three valine residues, V309, V350, and V363, display resolved Cγ chemical shifts for the two methyl groups. Cα-Cβ of alanine residues exhibit random coil chemical shifts as well. (b) Both R349 (located in R4) and Y310 (located in R3) have well-resolved Cζ signals at 159.6 ppm and 157.6 ppm, respectively.

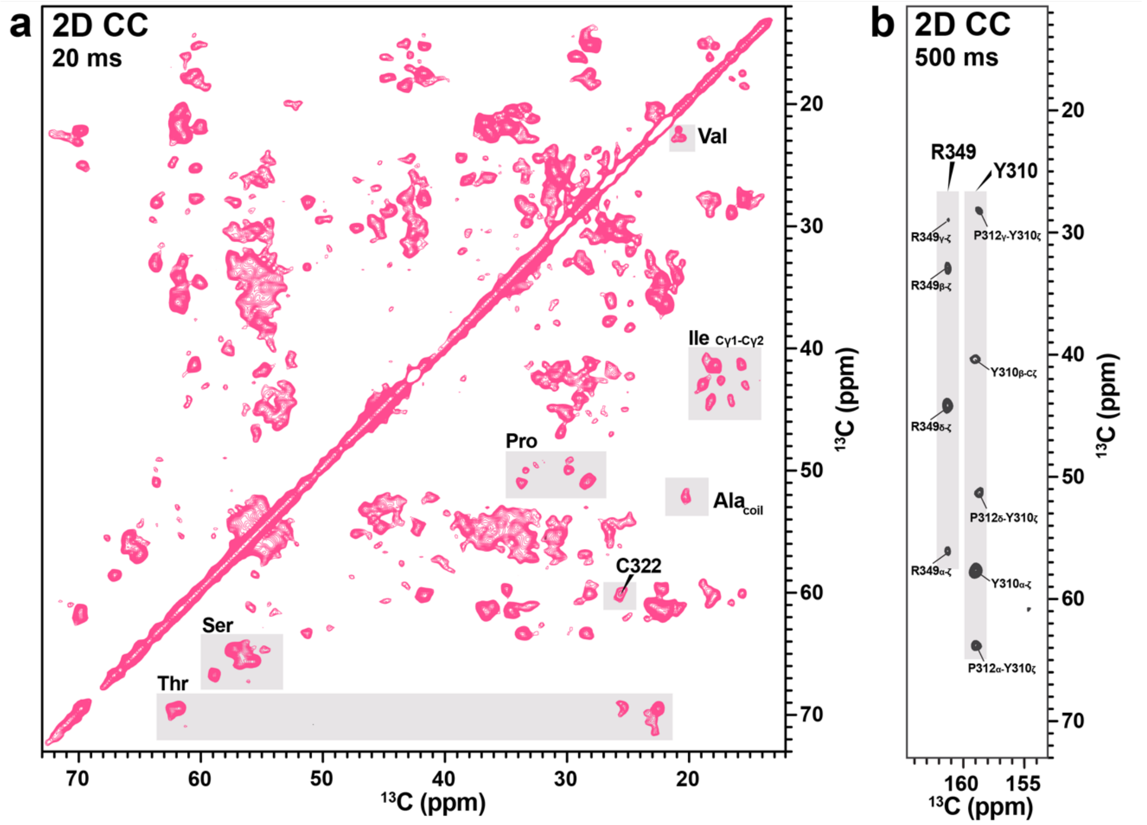

**Figure S3.** Backbone assignment of K321-L325 fragment in the dGAE filaments. Representative strip plot of NCACX (black), CANCO (blue), and NCOCX (red) spectra.

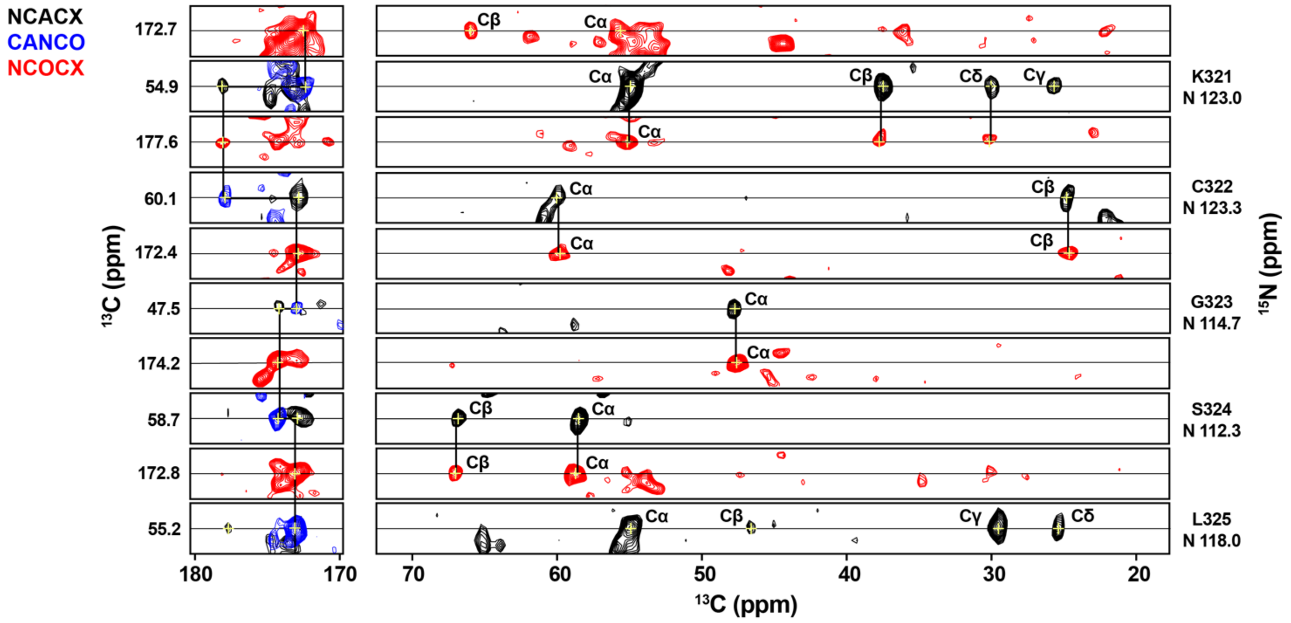

**Figure S4.** 2D NCA spectrum of selectively labeled dGAE filaments. (a) Lysine suppressed sample. (b) Valine, isoleucine, leucine, and lysine suppressed sample.

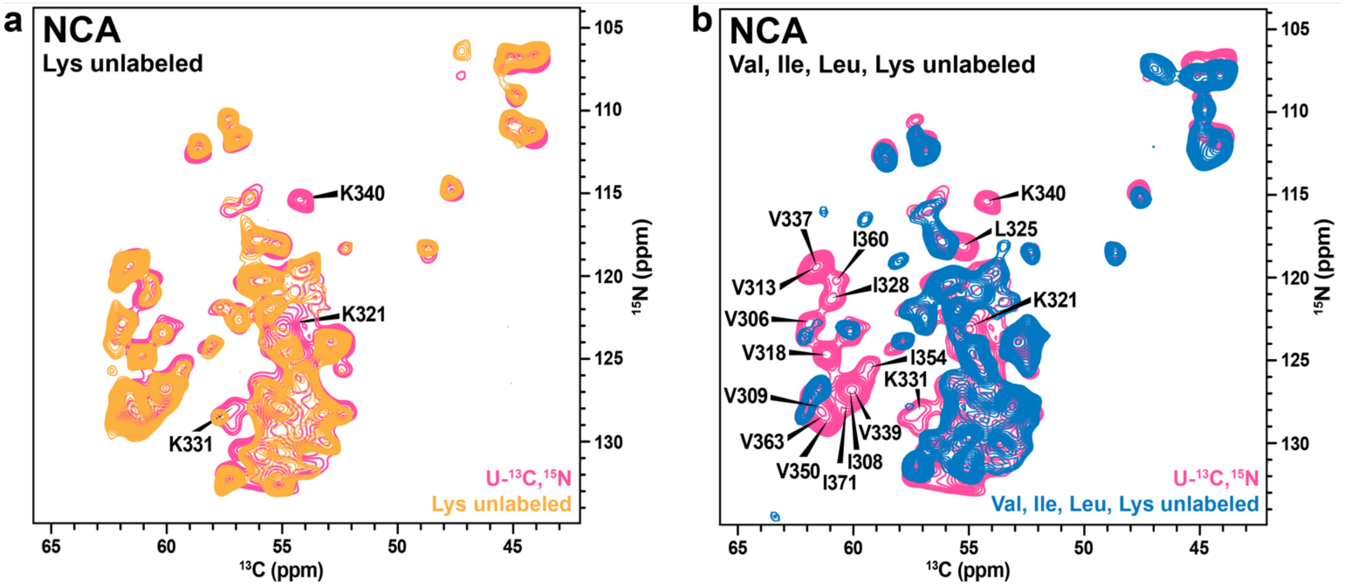

**Figure S5.** (a) Assigned 2D NCA spectrum of dGAE filaments. (b) 2D $^1$H-$^{13}$C INEPT spectrum of dGAE filaments.

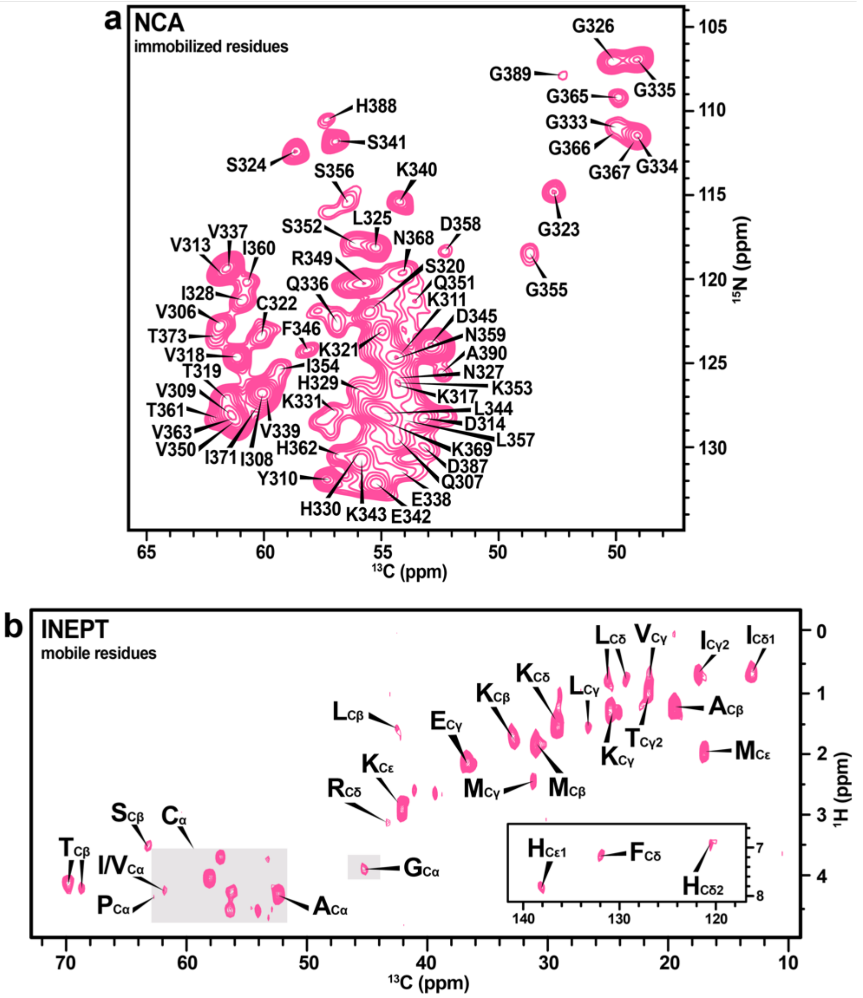

**Figure S6.** AFM image of the seeded full-length 2N4R tau filaments.

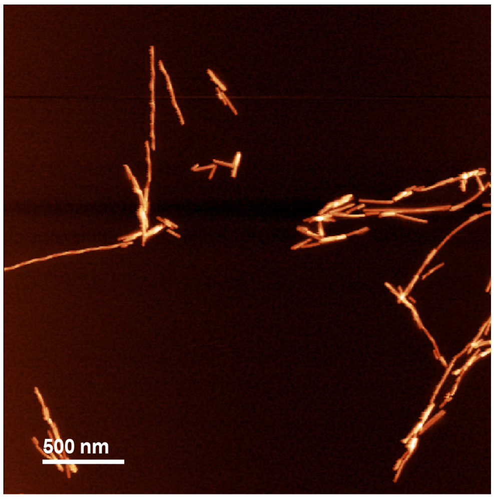

**Figure S7.** Overlay of 2D $^{1}H$-$^{13}C$ INEPT spectra of dGAE and seeded full-length tau filaments.

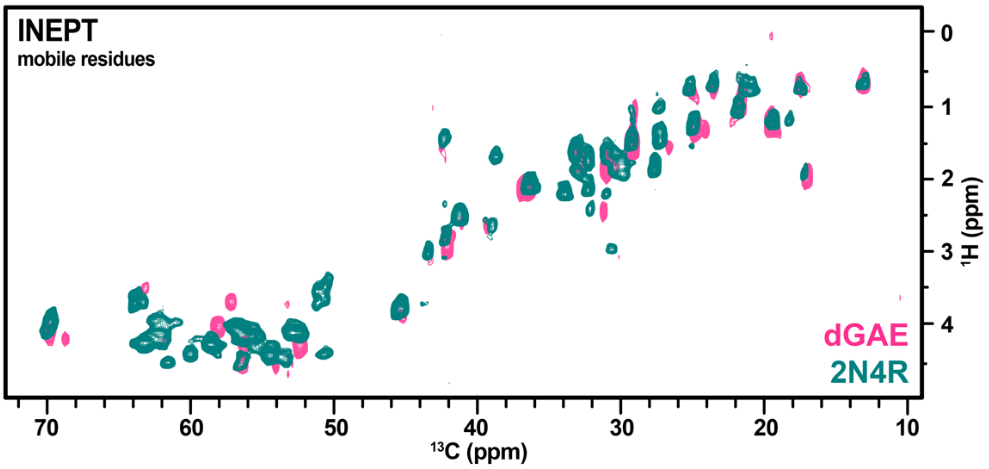

**Figure S8.** An amino acid sequence of dGAE with repeats highlighted.

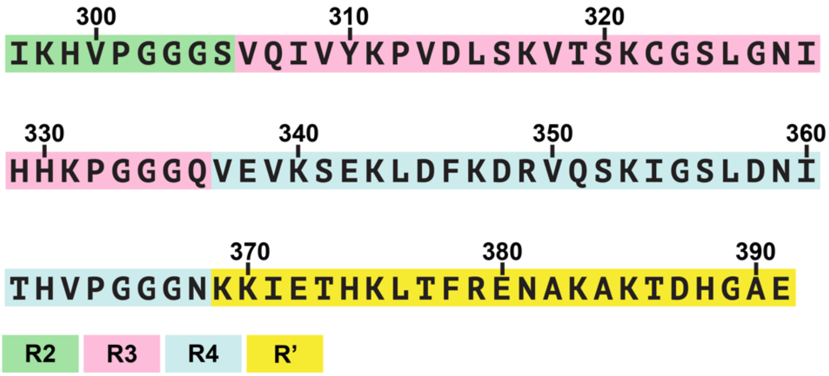

**Table S1.** Acquisition and processing parameters for the $^{13}C$-detected 800 MHz NMR spectra of uniformly labeled dGAE sample (U-$^{13}C$,$^{15}N$-dGAE), lysine unlabeled sample (noK-dGAE), and valine, isoleucine, leucine, and lysine unlabeled sample (noKLIV-dGAE).

| Sample | Spectrum | Dec. field ampl. (kHz) | CP conditions | Rec. delay (s) | Scans | Exp. time | Window function |
|---|---|---|---|---|---|---|---|
| U-$^{13}C$,$^{15}N$-dGAE | 2D $^{13}C$-$^{13}C$ DARR (20 ms mixing) | 100 | $^{1}H\rightarrow^{13}C$ 0.9 ms 59/47 kHz | 2.5 | 32 | 18 h 18 min | 60° shifted squared sine-bell |
| | 2D $^{13}C$-$^{13}C$ DARR (150 ms mixing) | 100 | $^{1}H\rightarrow^{13}C$ 0.9 ms 59/47 kHz | 2.5 | 160 | 60 h | 72° shifted squared sine-bell |
| | 2D NCA | 100 | $^{1}H\rightarrow^{15}N$ 1 ms 59/71 kHz<br>$^{15}N\rightarrow^{13}C$ 5.4 ms 47/59 kHz | 2.5 | 256 | 16 h | 72° shifted squared sine-bell |
| | 2D NCO | 100 | $^{1}H\rightarrow^{15}N$ 1 ms 59/71 kHz<br>$^{15}N\rightarrow^{13}C$ 6 ms 47/59 kHz | 3 | 64 | 5 h | 60° shifted squared sine-bell |
| | 2D CaN | 100 | $^{1}H\rightarrow^{13}C$ 0.9 ms 59/47 kHz<br>$^{13}C\rightarrow^{15}N$ 4.8 ms 47/59 kHz | 3 | 320 | 34 h 30 min | 60° shifted squared sine-bell |
| | 2D CoN | 100 | $^{1}H\rightarrow^{15}N$ 0.7 ms 59/47 kHz<br>$^{13}C\rightarrow^{15}N$ 4.2 ms 47/59 kHz | 3 | 268 | 41 h | 60° shifted squared sine-bell |
| | 2D NCACX | 100 | $^{1}H\rightarrow^{15}N$ 1 ms 59/71 kHz<br>$^{15}N\rightarrow^{13}C$ 5.4 ms 47/59 kHz | 3 | 256 | 17 h 30 min | 60° shifted squared sine-bell |
| | 3D NCACX | 100 | $^{1}H\rightarrow^{15}N$ 1 ms 59/71 kHz<br>$^{15}N\rightarrow^{13}C$ 5.4 ms 47/59 kHz | 2.5 | 128 | 234 h | 72° shifted squared sine-bell |
| | 3D NCOCX | 100 | $^{1}H\rightarrow^{15}N$ 1.1 ms 59/71 kHz<br>$^{15}N\rightarrow^{13}C$ 7 ms 47/59 kHz | 3 | 128 | 174 h | 60° shifted squared sine-bell |
| | 3D CANCO | 100 | $^{1}H\rightarrow^{13}C$ 1 ms 69/47 kHz<br>$^{13}C\rightarrow^{15}N$ 5.6 ms 59/47 kHz<br>$^{15}N\rightarrow^{13}C$ 5.2 ms 47/59 kHz | 3 | 256 | 262 h | 60° shifted squared sine-bell |
| | 2D $^{1}H$-$^{13}C$ INEPT | 100 | – | 2.5 | 64 | 11 h 30 min | 72° shifted squared sine-bell |
| noK-dGAE | 2D $^{13}C$-$^{13}C$ DARR (20 ms mixing) | 100 | $^{1}H\rightarrow^{13}C$ 0.9 ms 59/47 kHz | 2.5 | 32 | 18 h 18 min | 60° shifted squared sine-bell |
| | 2D $^{13}C$-$^{13}C$ DARR (500 ms mixing) | 100 | $^{1}H\rightarrow^{13}C$ 0.9 ms 59/47 kHz | 2.9 | 128 | 49 h | 60° shifted squared sine-bell |
| | NCA | 100 | $^{1}H\rightarrow^{15}N$ 1 ms 59/71 kHz<br>$^{15}N\rightarrow^{13}C$ 5.4 ms 47/59 kHz | 2.5 | 256 | 16 h | 60° shifted squared sine-bell |
| | NCO | 100 | $^{1}H\rightarrow^{15}N$ 1 ms 59/71 kHz<br>$^{15}N\rightarrow^{13}C$ 5.4 ms 47/59 kHz | 3 | 64 | 5 h | 60° shifted squared sine-bell |
| | 2D NCACX | 100 | $^{1}H\rightarrow^{15}N$ 1 ms 59/71 kHz<br>$^{15}N\rightarrow^{13}C$ 5.4 ms 47/59 kHz | 3 | 768 | 52 h | 60° shifted squared sine-bell |
| | 3D NCOCX | 100 | $^{1}H\rightarrow^{15}N$ 1 ms 59/71 kHz<br>$^{15}N\rightarrow^{13}C$ 7 ms 47/59 kHz | 2.8 | 64 | 59 h 30 min | 90° shifted squared sine-bell |
| noKLIV-dGAE | 2D $^{13}C$-$^{13}C$ DARR (20 ms mixing) | 100 | $^{1}H\rightarrow^{13}C$ 0.9 ms 59/47 kHz | 2.5 | 32 | 18 h 18 min | 60° shifted squared sine-bell |
| | 2D $^{13}C$-$^{13}C$ DARR (100 ms mixing) | 100 | $^{1}H\rightarrow^{13}C$ 0.9 ms 59/47 kHz | 2.7 | 256 | 101 h | 60° shifted squared sine-bell |
| | 2D $^{13}C$-$^{13}C$ DARR (500 ms mixing) | 100 | $^{1}H\rightarrow^{13}C$ 0.9 ms 59/47 kHz | 2.8 | 192 | 85 h | 60° shifted squared sine-bell |
| | NCA | 100 | $^{1}H\rightarrow^{15}N$ 1 ms 59/71 kHz<br>$^{15}N\rightarrow^{13}C$ 5.4 ms 47/59 kHz | 2.5 | 256 | 11 h 30 min | 60° shifted squared sine-bell |
| | NCO | 100 | $^{1}H\rightarrow^{15}N$ 1 ms 59/71 kHz<br>$^{15}N\rightarrow^{13}C$ 5.4 ms 47/59 kHz | 2.5 | 128 | 6 h | 60° shifted squared sine-bell |
| | 2D NCACX | 100 | $^{1}H\rightarrow^{15}N$ 1 ms 59/71 kHz<br>$^{15}N\rightarrow^{13}C$ 5.4 ms 47/59 kHz | 3 | 384 | 26 h | 60° shifted squared sine-bell |

**Table S2.** Acquisition and processing parameters for the $^{13}C$-detected 800 MHz NMR spectra of dGAE seeded full-length 2N4R tau sample.

| Sample | Spectrum | Dec. field ampl. (kHz) | CP conditions | Rec. delay (s) | Scans | Exp. time | Window function |
|---|---|---|---|---|---|---|---|
| **2N4R** | 2D $^{13}C$-$^{13}C$ DARR (20 ms mixing) | 100 | $^{1}H \rightarrow ^{13}C$ 0.6 ms 59/47 kHz | 2.7 | 64 | 34 h | 60° shifted squared sine-bell |
| | 2D NCA | 100 | $^{1}H \rightarrow ^{15}N$ 1.2 ms 59/71 kHz<br>$^{15}N \rightarrow ^{13}C$ 5.6 ms 47/59 kHz | 2.5 | 384 | 24 h | 60° shifted squared sine-bell |
| | 2D $^{1}H$-$^{13}C$ INEPT | 100 | – | 3 | 32 | 13 h 30 min | 60° shifted squared sine-bell |


### Author Contributions

K.J., R.S., J.H., and K.K. conceptualized and designed the project. K.J. supervised the project. K.K., A.L.B. carried out protein expression and purification with R.S help. K.K. designed and performed filament assembly. G. K. measured the EM and AFM images. K.K., L.K. carried out ThT experiments with V.S. help. A.L. recorded ssNMR experiments. K.K. assigned the NMR spectra with K.J. help. M.Z., B. K., R. S. performed proteophatic seeding activity experiments. All authors discussed the results. K.K. and K.J. wrote the manuscript with input from all authors. All authors read and approved the final version of manuscript.
